# Changes in bioactive components of *Aristolochia tagala*. Cham, a rare species of medicinal importance during its *in vitro* development through direct regeneration

**DOI:** 10.1101/037028

**Authors:** M Remya, V Narmatha Bai, S Murugesan, VN Mutharaian

## Abstract

Tissue culture propagation system was developed for *Aristolochia tagala*, a threatened medicinal plant, using apical bud explants. The most effective medium was found to be MS medium supplemented with BAP (3 μM), KIN (0.5 μM) and activated charcoal (0.1%). The addition of activated charcoal helped in circumventing the problem of polyphenol exudation from the explants which hampered the regeneration of adventitious shoots. A maximum of 12.6 shoots were obtained on average from the apical bud explants after 25 days of inoculation. Well developed shoots were rooted on MS medium supplemented with indole acetic acid (1.5 μM), Kinetin (1.5 μM) and 6-benzylaminopurine (0.5 μM). Regenerated shoots from the apical buds were successfully rooted and acclimated to greenhouse conditions. Qualitative and quantitative analysis of bioactive compounds was done at various stages of development, so as establish the effect of culture conditions on the production of bioactive components. Comparisons were made between three types of plant material from the same clone: leaves from field-grown plant, leaves from *in vitro* apical bud cultures and leaf derived callus. It was observed that the leaf derived callus showed the presence of components which were not there in the *in vivo* leaves, suggesting the influence of *in vitro* developmental conditions.

**Tagala:** Summary Statement

This is the first report on direct regeneration of *Aristolochia tagala*, (a threatened yet important medicinal plant) using apical bud explants derived from mature plants. Also, for the very first time we report the results of analysis of secondary metabolites during its various stages of *in vitro* development.

**Abbreviations:** AC - activated charcoal; BAP – 6-benzylaminopurine; 2,4-D - 2,4- dichloro phenoxyacetic acid, IAA - indole acetic acid, IBA - indolebutyric acid, KIN- kinetin, MS- Murashige and Skoog (1962); NAA- α-naphthaleneacetic acid.

## Introduction

*Aristolochia* L. is a large genus of Aristolochiaceae with about 120 species, distributed throughout tropical and subtropical countries. *Aristolochia* species have been studied for their pollination strategies associated with floral aroma and fly-trapping perianth, as well as for their species-specific host plant relationships with swallowtail butterfly larvae (Banziger and Disney, 2006; Murugan et al., 2006; Trujillo and Sersic, 2006). Secondary metabolites produced by *Aristolochia* plants are critical for the defense and survival of the butterflies during their larval feeding stage, such that the decline of particular swallowtail butterfly population is attributed to the declining distribution of particular *Aristolochia* species (Klitzke ad Brown 2000; Kumar et al., 2003).

*Aristolochia tagala* Cham., a climbing shrub is distributed in India, Sri Lanka, China, Malaysia, Burma, Java and Australia, and is a rare medicinal plant, Murugan et al., (2006).Various parts of *A. tagala* are extensively used in traditional medicine. The juice from the leaves is used as a specific antidote for Cobra poison, Kirtikar and Basu (2003). The roots are strongly aromatic and are used to treat snake bites, bone fractures, malaria, indigestion, rheumatism, toothaches and various dermatological conditions by the Kani tribe of Thiruvananthapuram and Tirunelveli hills, India. Roots are also used for a medicated steam bath, known as ‘sudorification’. Leaves are used to treat colic fits and bowel complaints. Due to indiscriminate harvesting of roots for local medicine and trade, the species has become rare in its natural habitat, Ravikumar and Ved (2000).

Propagation of *A. tagala* relies solely on seeds. However the viability of the seeds is very low due to the presence of scanty endosperm, Biswas et al., (2007). This calls for newer approaches to the rapid propagation of this important medicinal plant. Plant tissue culture techniques are now being used globally for the conservation, and multiplication of medicinally important plant species. Micropropagation holds promise as a major component in medicinal plant breeding within a reasonable time frame without affecting the wild bioresources.

To the best of our knowledge, this is the first report on regeneration of *A. tagala* using apical bud explants derived from mature plants. Also, there are no reports of analysis of secondary metabolites from the *in vitro* derived plantlets of this medicinally important species. Therefore the present study is aimed at testing the efficiency of apical bud explants to regenerate into *A.tagala* plantlets and also at exploring the bioactive compounds present during the different stages of *in vitro* development.

## Materials and Methods

### Plant material and source of explant

*A. tagala* was collected from Kolli hills, which is situated in the south western edge of the Eastern Ghats, India. The plant was identified by Dr. R. Gopalan (Botanical Survey of India, Southern Circle, Coimbatore, India), and a voucher specimen of the same was deposited in the herbarium of the Botanical Survey of India (Accession number-167986). To avoid causalities, some of the plants were maintained under green house conditions at the Department of Botany, Bharathiar University, Coimbatore, India.

### Explants selection and mode of sterilization

Shoot tips (5-10 mm) from actively growing plants were collected and and washed thoroughly in running tap water for 20- 30 min followed by treatment with 5% (v/v) Teepol (detergent) for 5 – 10 min. The treated explants were washed with distilled water for 5 min. Subsequently the explants were disinfected with 0.1% mercuric chloride for 3-5 min and rinsed 3- 4 times with sterile distilled water under aseptic conditions inside the laminar hood. Apical buds excised from the surface sterilized shoot tips with the help of sterile surgical blade (Lisyter, No. 10) were used as the explants.

### Direct shoot regeneration from the apical buds

The explants were aseptically cultured on regeneration medium, Murashige and Skoog (1962). The regeneration medium was supplemented with 3% (W/V) sucrose and consisted of full strength MS macro nutrients, micro nutrients and vitamins. The pH of the regeneration medium was adjusted to 5.8 before adding 0.8% (W/V) agar (Hi Media, Mumbai). 15 cm^3^ of the medium was poured into 25X150 mm culture tubes (Borosil, Mumbai) and autoclaved at 121°C and 1.06 kg cm^−2^ pressure for 20 min. Two explants were inoculated per tube. The cultures were incubated at 25 ± 2°C under a 16h photoperiod of 50-60 μmol m^−2^ s^−1^ flux density provided by cool white fluorescent tubes (Philips).

MS medium supplemented with different combinations of BAP (0.5 – 3 μM), KIN (0.5 – 2 μM), 2,4-D ( 0.5 - 1μM) and 0.1% activated charcoal (w/v) was used to find an optimum culture medium for inducing direct adventitious shoot regeneration. The time taken for the induction of multiple shoots, total number of shoots per explant and the length of the shoots were recorded.

### Elongation and rooting of the micro shoots

The micro shoots obtained from the apical bud explants were sub cultured in MS solid and liquid medium supplemented with GA_3_ (0.5 – 2 μM) to facilitate early elongation. The elongated micro shoots (10- 14 cm) were transferred to MS medium supplemented with growth regulators like IAA, KIN, BAP, individually and in combinations at concentrations ranging from 0.5 – 1.5mg l^−1^. The cultures were kept in the dark at 25 ± 2°C. Data were recorded on length of the roots after 3 weeks of transfer onto the rooting media.

### Hardening

Acclimatization and hardening of the *in vitro* developed plantlets were carried out using the protocol standardized by us earlier (Remya et al., 2013). Well-developed, healthy plantlets with approximately 6 cm long shoots (containing 6 -10 leaves ranging from 0.5 – 1 cm) and approximately 13 cm long tap roots were removed from the culture flasks and were washed thoroughly in running tap water to remove the adhering nutrient medium. The plantlets were then soaked in 1% (w/v) of the fungicide, methyl-3-benzimidizole carbomate (Bavistin) for 10-15 min and transferred to small plastic pots filled with various types of sterilized potting mixtures.

The plantlets were initially maintained under laboratory conditions at 25 ± 2°C for 10 d. During this period they were sprayed with liquid MS medium (quarter strength-without sucrose) daily and covered tightly with polyethylene bags. After 10 d they were gradually exposed to the natural environmental conditions.

### Callus induction from leaf explants

The surface sterilized leaf explants were trimmed gently with the help of a sterile surgical blade (Lisyter, No. 10) and aseptically cultured on regeneration medium (Murashige and Skoog 1962).The regeneration medium was supplemented with 3% (W/V) sucrose and consisted of full strength MS macro nutrients, micro nutrients and vitamins. The pH of the regeneration medium was adjusted to 5.8 before adding 0.8% (W/V) agar (Hi Media, Mumbai). 15 cm^3^ of the medium was poured into 25X150 mm culture tubes (Borosil, Mumbai) and autoclaved at 121°C and 1.06 kg cm^−2^ pressure for 20 min. Two explants were inoculated per tube. The cultures were incubated at 25± 2°C under a 16h photoperiod of 50-60 μmol m^−2^ s^−1^ flux density provided by cool white fluorescent tubes (Philips).

MS medium enriched with auxins (2, 4-D & NAA) and cytokinins (BAP & KIN) at concentrations ranging from 0.5 – 3.0 μM was used for callus induction. The calli were sub cultured regularly at an interval of four weeks.

### Analysis of bioactive compounds

#### a) Sugars (Harborne, 1960)

The plant tissues namely *in vivo* leaves, leaf derived callus and *in vitro* leaves of *A. tagala* were dried, powdered and extracted with 95% methanol. The extract was concentrated to remove methanol and filtered through Whatman No.1 filter paper. The clear extract was spotted directly on the TLC plates.

Total sugars were quantified by the Dubois (1956) method

#### b) Alkaloids (Harborne, 1973 a)

The alkaloids were extracted from the plant tissues using 1N HCl and the crude extract was left for 12 hrs at room temperature. The crude extract was filtered and concentrated to one quarter of the original volume. To the above extract, concentrated ammonia solution was added drop by drop for precipitation and centrifuged with 1N ammonia solution and then the pellets were spotted on TLC plates. The bands formed on the TLC plates were scraped out and and the compounds were identified using UV Visible spectrophotometer (Shimadzu 1601) at the range of 200-700 nm using methanol as the blank.

Total alkaloid content was determined by the method of Fazel et al., 2008. The plant extract was dissolved in 2 N HCl and then filtered. The pH of phosphate buffer solution was adjusted to neutral with 0.1 N NaOH. One ml of this solution was transferred to a separating funnel and then 5 ml of bromocresol green (BCG) solution along with 5 ml of phosphate buffer were added. The mixture was shaken and the complex formed was extracted with chloroform by vigorous shaking. The extracts were collected in a 10 ml volumetric flask and diluted to volume with chloroform. The absorbance of the complex in chloroform was measured at 470 nm.

#### c) Phenols (Harborne, 1973 b)

The dried powder of various plant tissues (10 gram each) was hydrolyzed with 2N HCl at 40°C for half an hour. It was allowed to cool at room temperature and filtered. The filtrate was washed with equal volume of petroleum ether using a separating funnel. The extracts so obtained were spotted on the TLC plates. The bands so formed were scraped out and and the compounds were identified using UV Visible spectrophotometer (Shimadzu 1601) at the range of 200-700 nm using methanol as the blank.

Folin Ciocalteu method was used to determine the total phenolic contents of methanolic extract by UV spectrophotometer. An external calibration curve of gallic acid as standard phenolic compound was plotted. The total phenolic contents of the methanolic extract of fruits were calculated with the help of a standard calibration curve[Mc Donald], and are reported as gallic acid equivalent (mg/g of dry mass).

#### d) Flavonoids (Bate-Smith, 1962)

10 gm of the different samples were immersed in 2N HCl and heated for 30-40 min at 100°C. The extracts were then cooled at room temperature and the cooled extracts were washed with ethyl acetate using a separating funnel. The ethyl acetate extracts were concentrated to dryness, taken up in 1-2 drops of ethanol and aliquots were chromatographed one dimensionally.

A small spot was applied to the silica gel plates and the chromatograms were developed in various solvent systems as shown in Table-1. The bands so formed were scraped out and and the compounds were identified using UV Visible spectrophotometer (Shimadzu 1601) at the range of 200-700 nm using methanol as the blank.

**Table- 1.** Different solvent systems used in the TLC

| Compounds | Solvent system used | Ratio |
| --- | --- | --- |
| Sugar | Butanol: Acetic acid: Distilled water<br>(The top layer was used) | 4:1:5 |
|  | Benzene: Ethyl acetate: Distilled water<br>(The top layer was used) | 4:1:2 |
| Alkaloids | Methanol: Conc. Ammonium Hydroxide | 200:5 |
| Phenols | Acetic acid: Chloroform | 1:9 |
|  | Ethyl acetate: Benzene | 9:11 |
|  | Benzene: Methanol: Acetic acid | 45:8:4 |
| Flavonoids | Acetic acid: HCl: Distilled water<br>(The top layer was used) | 10:3:30 |
|  | Butanol: Acetic acid: Distilled water<br>(The top layer was used) | 4:1:5 |

Aluminum chloride colorimetric method of Chang et al., 2002 was used to analyze the flavonoid content. Quercetin (standard flavonoid compound) was used to construct a standard calibration curve. The flavonoid contents in the extract is reported as quercetin equivalent (mg/g of dry mass).

Analysis of bio active components using GC/MS:

1μ1 of the methanolic extracts of leaves from field-grown plant, *in vitro* apical bud cultures and leaf derived callus of *A. tagala* were injected in GC-MSD (Fissions 8000 series Gas chromatograph with Fisons 800 series Mass spectrophotometer). The column used was DB-5 MS and the flow rate was maintained at 1.2 ml/min with helium as the carrier gas. The temperature of the injector and detector was 275 °C. The oven temperature was programmed from 35 to 200°C at a rate of 4°C /min. Identification of components was based on their retention indices, which were determined with references to a series of standards, and by comparison of their mass spectral fragmentation pattern with NIST database (John Wiley Library 229119), Adams (2001).

#### Statistical analysis

All the experiments were conducted with a minimum of 10 replicates per treatment. The experiments were repeated three times. The significance of differences among means was carried out using Duncan’s multiple range test (DMRT) at P = 0.05. The results are expressed as means ± SE of three experiments.

### Results

#### Induction of adventitious shoots

Apical bud explants cultured on MS medium fortified with BAP (0.5 μM) produced only one or two shoots. It was observed that there was polyphenol exudation from the explants which hampered the regeneration of adventitious shoots. Therefore, we tried the addition of activated charcoal to the regeneration medium. When activated charcoal (0.1%) was added to MS medium supplemented with growth hormones, it evoked shoot bud regeneration. It was observed that a combination of BAP (3 μΜ), KIN (0.5 μΜ) and AC (0.1%), induced the maximum number of shoots (Tab. 2) (Fig. 1a).

**Table-2.** Effect of growth regulators and activated charcoal on the production of multiple shoots from the apical buds of *A. tagala*.

| BAP | KIN | 2,4-D | AC | No. of shoots | Length of the shoots | Time taken |
| --- | --- | --- | --- | --- | --- | --- |
| (μM) | (μM) | (μM) | (%) | | (cm) | (d) |
| 0.5 | - | - | 0.1 | 1.20± 0.12<sup>a</sup> | 1.42 ± 0.26<sup>a</sup> | 28± 0.23<sup>g</sup> |
| 0.5 | 1 | - | 0.1 | 3.80 ± 0.16<sup>bc</sup> | 1.52 ± 0.12<sup>a</sup> | 20± 0.13<sup>f</sup> |
| 0.5 | 1.5 | 1 | 0.1 | 3.0± 0.22<sup>b</sup> | 1.46± 0.17<sup>a</sup> | 33± 0.22<sup>g</sup> |
| 0.5 | 2 | 0.5 | 0.1 | 5.60± 0.18<sup>de</sup> | 1.36 ± 0.34<sup>a</sup> | 30± 0.16<sup>i</sup> |
| 1 | 0.5 | - | 0.1 | 4.20± 0.23<sup>cd</sup> | 1.84± 0.22<sup>bc</sup> | 30± 0.24<sup>fi</sup> |
| 1 | 1 | - | 0.1 | 5.0± 0.25<sup>cd</sup> | 1.82± 0.21<sup>b</sup> | 30± 0.18<sup>d</sup> |
| 1 | 1.5 | - | 0.1 | 7.60± 0.11<sup>f</sup> | 2.04± 0.37<sup>cde</sup> | 28± 0.31<sup>g</sup> |
| 2 | 0.5 | - | 0.1 | 7.0 ± 0.26<sup>f</sup> | 2.30± 0.11<sup>f</sup> | 28± 0.28<sup>a</sup> |
| 2 | 1 | - | 0.1 | 9.20± 0.31<sup>f</sup> | 2.30 ± 0.35<sup>f</sup> | 28± 0.26<sup>f</sup> |
| 2 | 1.5 | - | 0.1 | 8.40± 0.18<sup>f</sup> | 1.98 ± 0.16<sup>bcd</sup> | 20± 0.23<sup>g</sup> |
| 3 | 0.5 | - | 0.1 | 12.60± 0.32<sup>f</sup> | 2.12± 0.36<sup>def</sup> | 20± 0.11<sup>i</sup> |
| 3 | 1 | - | 0.1 | 8.40± 0.14<sup>f</sup> | 2.22± 0.24<sup>cf</sup> | 20± 0.19<sup>f</sup> |
| 3 | 1.5 | - | 0.1 | 9.20± 0.28<sup>f</sup> | 2.60± 0.28<sup>g</sup> | 20± 0.25<sup>g</sup> |

Values are mean ± SE
In a column, means followed by a common letter are not significantly different at the 5% level by DMRT

**Fig:1.**
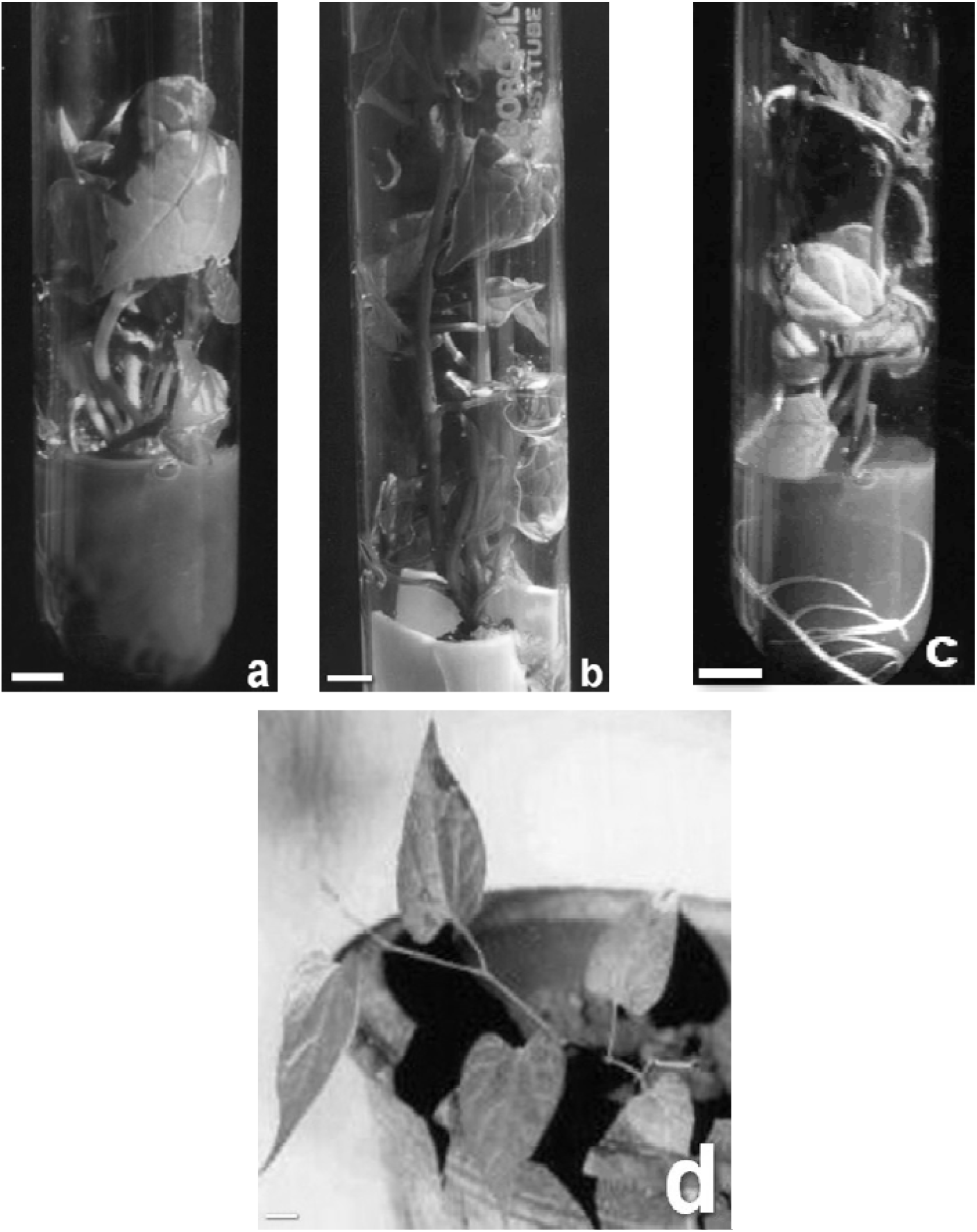
a. Multiple shoots from the apical bud explants of *A. tagala* b. Elongated micro shoots of *A. tagala* c. Rooted micro shoots of *A. tagala* d. Plantlet of *A. tagala*, growing in garden soil after hardening Bar = 1cm

#### Elongation of micro shoots

Elongation of micro shoots was achieved by sub culturing the micro shoots in MS medium supplemented with GA_3_. 2 μM was found to be the optimal concentration for shoot elongation. It was observed that MS liquid medium supplemented with GA_3_ was more effective when compared to solid medium (Tab. 3) (Fig.1b)

**Table -3.** Effect of GA _3_ on the elongation of micro shoots

In a column, means followed by a common letter are not significantly different at the 5% level by DMRT Table –3 Effect of GA<sub>3</sub> on the elongation of micro shoots
| Medium | GA<sub>3</sub><br>(μM) | Shoot length<br>(cm) | Time taken for elongation<br>(d) |
| --- | --- | --- | --- |
| MS + AGAR<br>(SOLID) | 0.5 | 6.6 ± 0.34<sup>a</sup> | 18 ± 0.11<sup>d</sup> |
|  | 1 | 9.78 ± 0.21<sup>b</sup> | 15 ± 0.13<sup>c</sup> |
|  | 2 | 14.8 ± 0.22<sup>d</sup> | 10 ± 0.21<sup>d</sup> |
| MS - AGAR<br>(LIQUID) | 0.5 | 8.98 ± 0.26<sup>b</sup> | 15 ± 0.23<sup>b</sup> |
|  | 1 | 12.86 ± 0.31<sup>c</sup> | 14 ± 0.25<sup>d</sup> |
|  | 2 | 14.96 ± 0.12<sup>d</sup> | 10 ± 0.15<sup>b</sup> |

Values are mean ± SE
In a column, means followed by a common letter are not significantly different at the 5% level by DMRT

#### Rooting of micro shoots

MS medium supplemented with IAA (1.5 μM), KIN (1.5 μM) and BAP (0.5 μΜ) was found to be the most effective in inducing roots. The elongated micro shoots when subcultured in the above mentioned medium produced roots of maximum length (18.4 cm) (Tab. 4) (Fig. 1c).

**Table-4.**
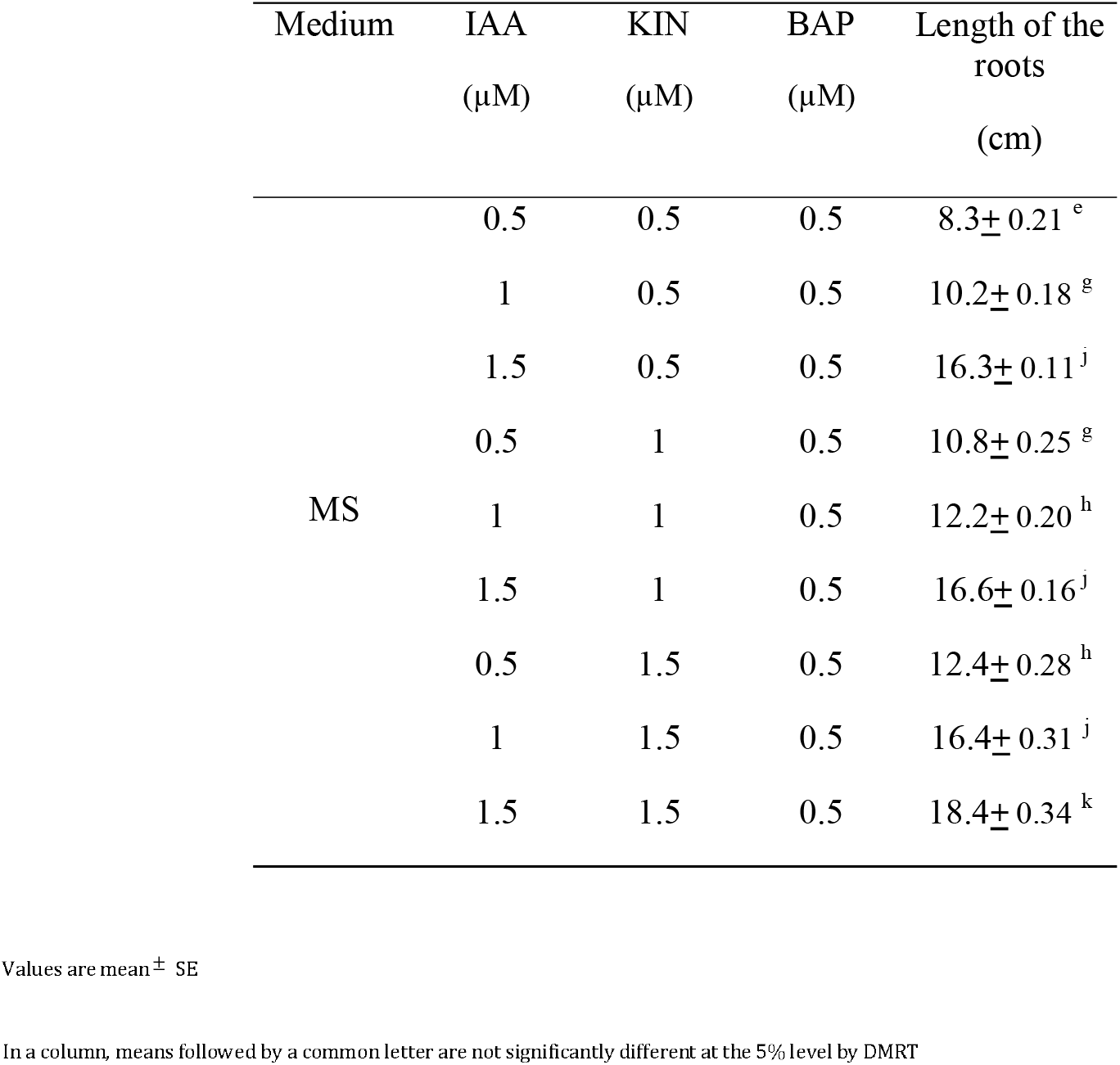
Effect of growth hormones on the induction of roots from the microshoots

#### Hardening

Plantlets with six to ten fully expanded leaves and well-developed roots obtained from the rooted micro shoots were successfully acclimatized in plastic pots containing coir pith along with vermiculite and soil in the ratio 1:1:1. The maximum percentage survival was recorded to be 80%. They were successfully transplanted into field conditions where they grew normally without any morphological variations. A well developed *in vitro* derived plantlet *A.tagala* is shown in Fig. 1d.

#### Callus induction

Best callusing (maximum dry weight of callus within minimum duration) was obtained when MS medium was fortified with BAP (1μM), NAA (0.5μM) and KIN (2μΜ).

#### Analysis of Bioactive compounds

The results of qualitative and quantitative analysis of the bioactive compounds from the leaves of field-grown plant, *in vitro* apical bud cultures and leaf derived callus of *A. tagala* are shown in Tab. 5–8. Flavonoids and Phenols were the dominant class of compounds in all the three extracts. It was observed that the *in vitro* leaves showed the presence of new components in addition to those which were expressed in the wild. The leaf derived calli also exhibited the presence of many new compounds which were neither present in the *in vivo* leaves, nor in the *in vitro* leaves.

**Table-5.** Qualitative and Quantitative analysis of alkaloids identified from the different samples of *A. tagala*

In a column, means followed by a common letter are not significantly different at the 5% level by DMRT Table-5 Qualitative and Quantitative analysis of alkaloids identified from the different samples of *A. tagala*
| Sample | Rf value<br>(x 100) | Absorption maxima<br>(nm) | Identified alkaloid | Amount of alkaloid<br>(mg/g ± S.E) |
| --- | --- | --- | --- | --- |
| *In vivo* leaves | 8 | 227 | Berberine | 22.45 ± .56 |
| *In vitro* leaves | 8 | 227 | Berberine | 24.32 ± .12 |
| Callus | - | - | - | - |

**Table 6.**
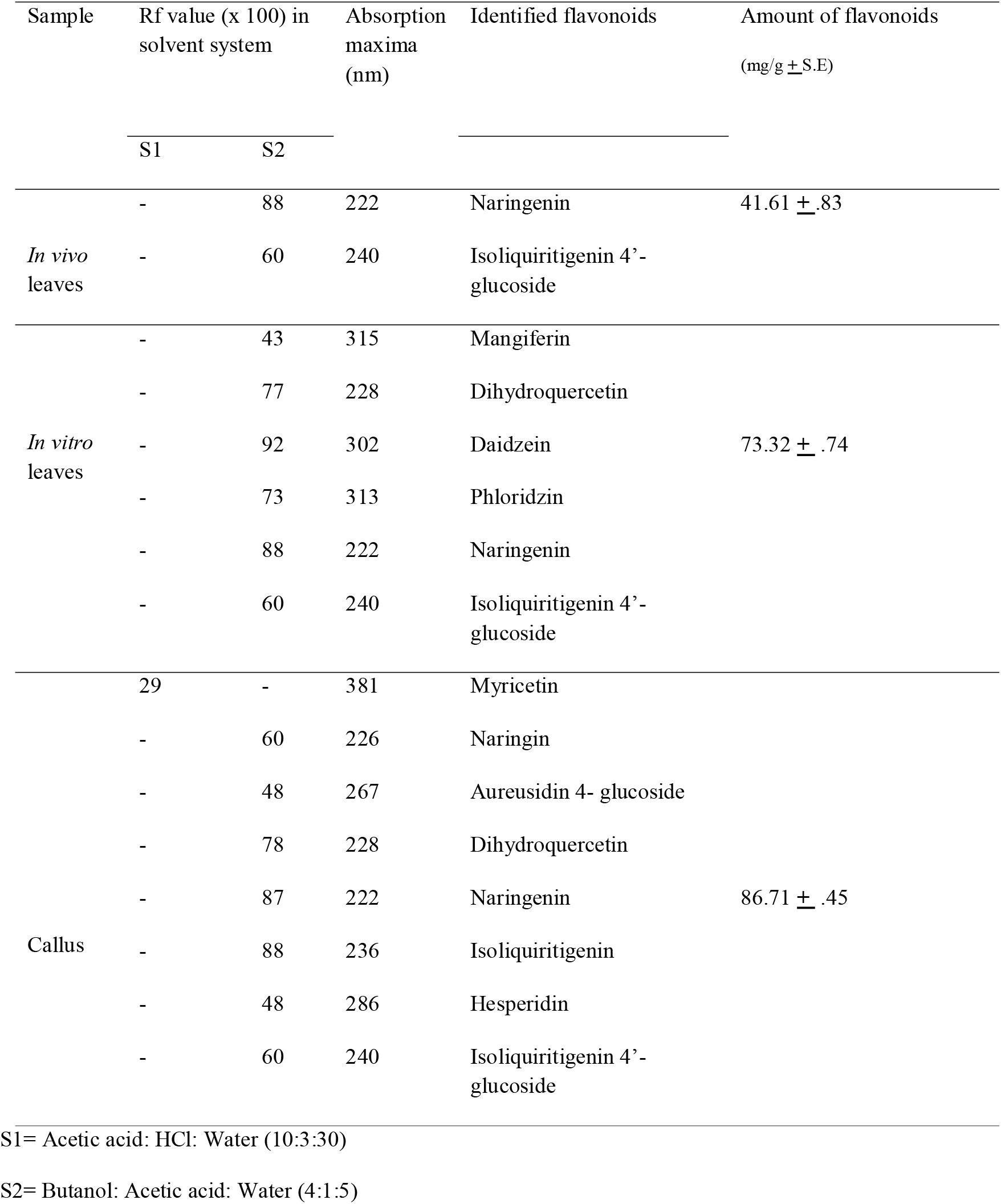
Qualitative and Quantitative analysis of flavonoids identified in the different samples of *A. tagala*

**Table 7.**
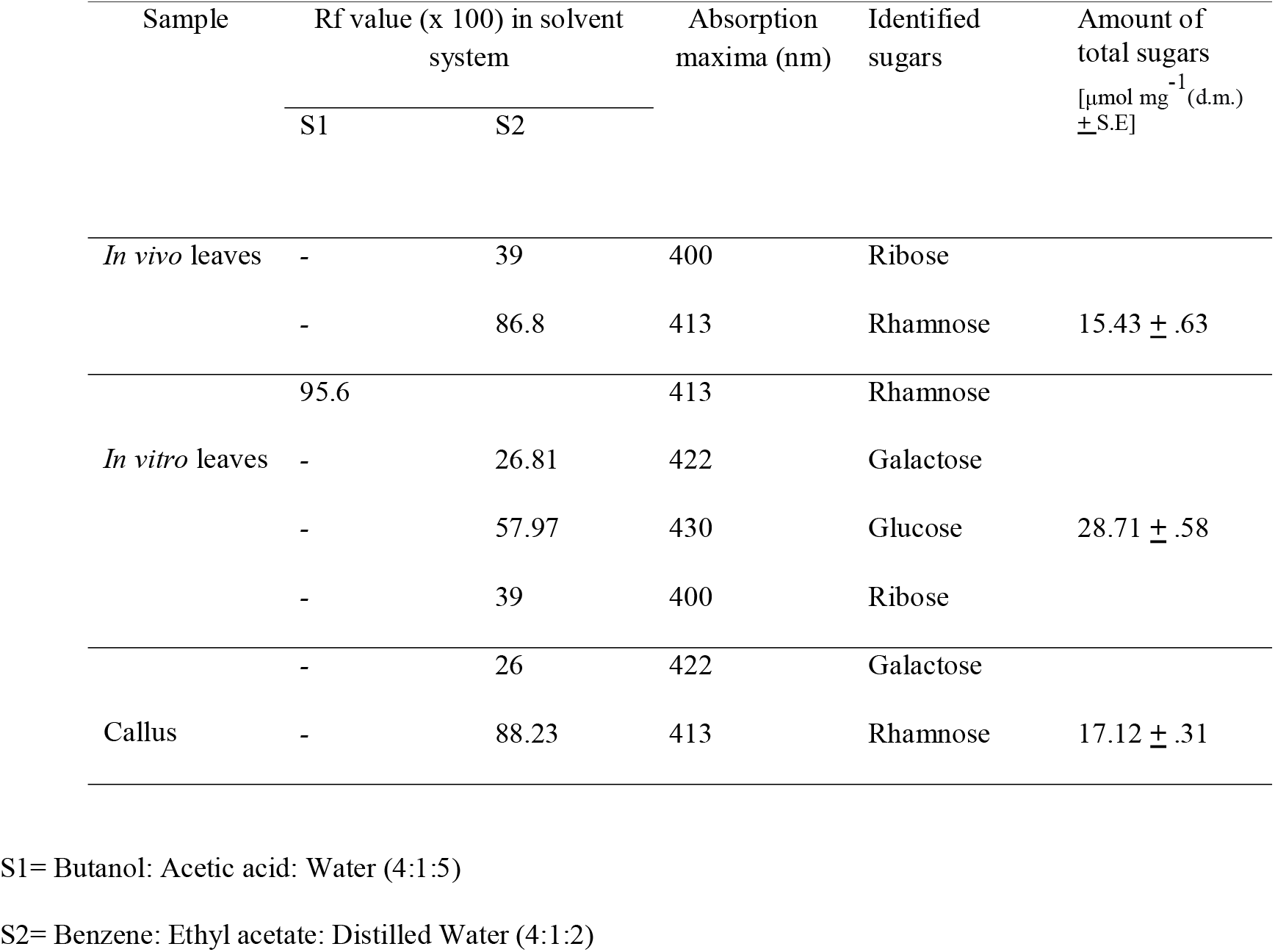
Qualitative and Quantitative analysis of sugars identified in the different samples of *A. tagala*

**Table-8.**
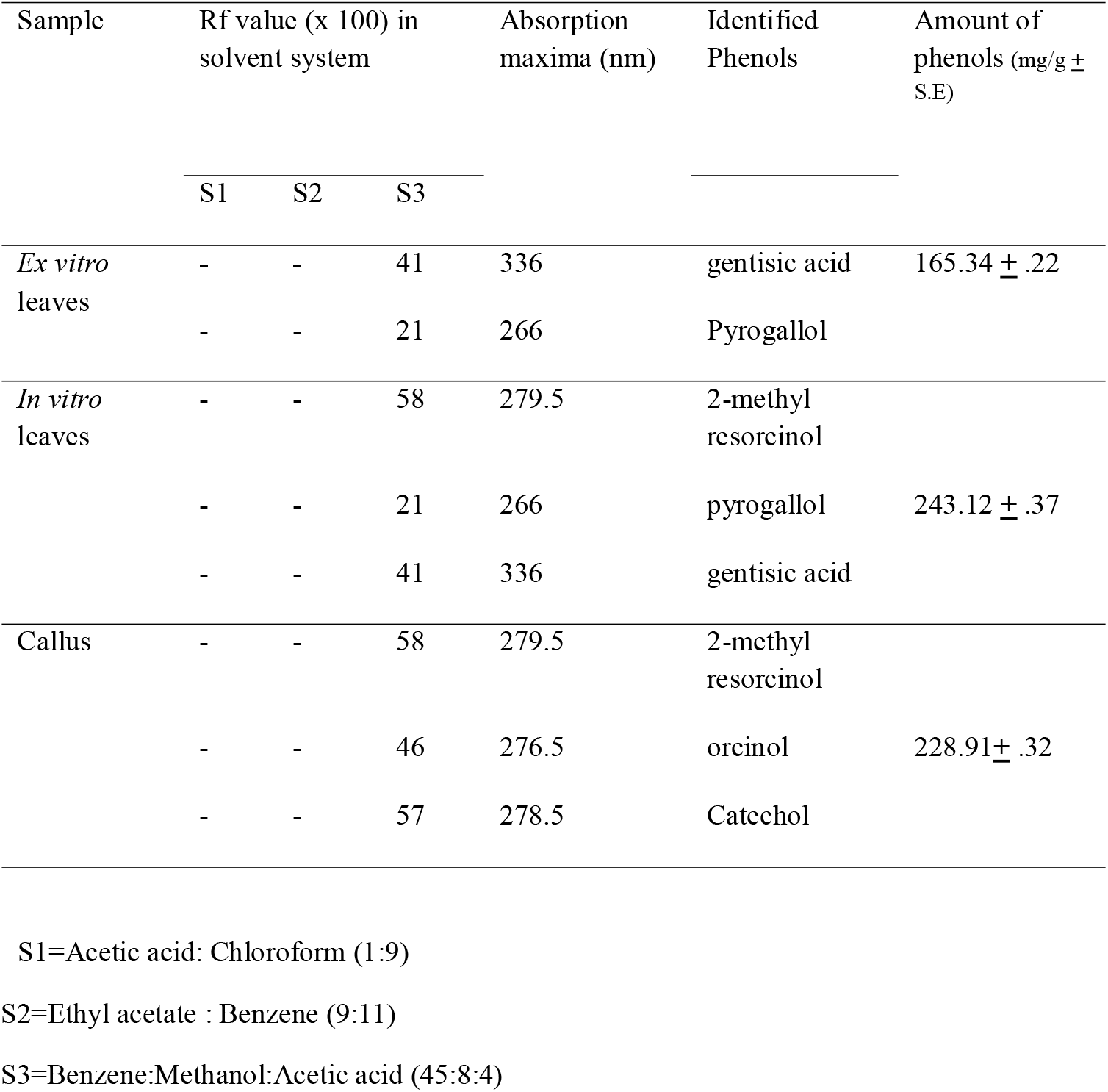
Qualitative and Quantitative analysis of phenols identified in the different samples of *A.tagala*

| Sample | Rf value (x 100) in solvent system | | | Absorption maxima (nm) | Identified Phenols | Amount of phenols (mg/g ± S.E) |
| --- | --- | --- | --- | --- | --- | --- |
|  | S1 | S2 | S3 |  |  |  |
| *Ex vitro* leaves | - | - | 41 | 336 | gentisic acid | 165.34 ± .22 |
|  | - | - | 21 | 266 | Pyrogallol |  |
| *In vitro* leaves | - | - | 58 | 279.5 | 2-methyl resorcinol |  |
| | - | - | 21 | 266 | pyrogallol | 243.12 ± .37 |
|  | - | - | 41 | 336 | gentisic acid |  |
| Callus | - | - | 58 | 279.5 | 2-methyl resorcinol |  |
| | - | - | 46 | 276.5 | orcinol | 228.91 ± .32 |
|  | - | - | 57 | 278.5 | Catechol |  |

S1=Acetic acid: Chloroform (1:9)
S2=Ethyl acetate : Benzene (9:11)
S3=Benzene:Methanol:Acetic acid (45:8:4)

GCMS analysis of the methanolic extracts of the *in vivo* leaves, *in vitro* leaves from apical bud cultures and leaf derived callus of *A.tagala*, revealed the presence of sesquiterpenes and higher fatty acids. A total of 20 compounds were identified from the *in* vivo and *in vitro* leaves (Tab. 9). It was observed that the same major components were found without significant compositional variations in the *in vivo* leaves and the *in vitro* leaves from direct regeneration.

**Table-9.** Constituents of the Methanolic extracts of the leaves of *A.tagala* identified by GCMS

| Compound No. | Name |
| --- | --- |
| 1 | Ethanone |
| 2 | 3,3-Dimethyl-5,6-Dihydroxy-Bicyclo[2,2,1]<br>Heptan-2-Carboxaldehyde |
| 3 | 4-Methyl-2,5-Dimethoxybenzaldehyde |
| 4 | 3-T-Butyl-\$-Hydroxyanisole |
| 5 | Phenol,(1,1-Dimethylethyl)-4-Methoxy |
| 6 | 3-Isopropyl-1,2-Dimethoxybenzene |
| 7 | Kongol |
| 8 | Junipene |
| 9 | 1-Naphthalenepropanol |
| 10 | 5 Acetoxymethyl-2,6,10-Trimethyl-2,9-Undecadien-6-Ol |
| 11 | Ishwarane |
| 12 | Delta-Guaiene |
| 13 | Palmitic acid |
| 14 | Capric acid |
| 15 | Lauric acid |
| 16 | Margaric Acid |
| 17 | Stearic acid |
| 18 | Myristic acid |
| 19 | Oleic Acid |
| 20 | Tridecanoic Acid |

In the present study, we report for the first time presence of a sesquiterpene hydrocarbon, Ishwarane, in the *in vivo* and *in vitro* leaves of *A.tagala*. The same compound was not present in the leaf derived callus of *A.tagala*. The leaf derived callus showed the presence of components which were not there in the *in vivo* leaves, suggesting the influence of culture conditions (Tab. 10)

**Table-10.** Constituents of the Methanolic extracts of the leaf derived callus of *A.tagala* identified by GCMS

| Compound No. | Name |
| --- | --- |
| 1 | Oxinol |
| 2 | Benzenamine |
| 3 | 1- Naphthalenamine 5,6,7,8-<br>Tetrahydron |
| 4 | 5-Methoxy 2-3 dimethylindole |
| 5 | 1-Naphthalenol 5,6,7,8- Tetrahydro- 2,5,<br>dimethyl |
| 6 | Indoline 3 ethyl 2,3 Dimethyl |
| 7 | Quinoline |
| 8 | Benzo 13 Thiopene,2,5,7-Trimethyl |
| 9 | Ethanone |
| 10 | Palmitic acid |
| 11 | Lauric acid |
| 12 | Margaric Acid |
| 13 | Stearic acid |
| 14 | Myristic acid |
| 15 | Octadecanoic acid |
| 16 | Tridecanoic Acid |

## Discussion

In Aristolochiaceae, *in vitro* differentiation of shoots as well as plantlets have been reported from a variety of explants such as leaves (Manjula et al., 1997), stem (Remashree et al., 1997), nodal segments (Remashree et al., 1994, Manjula et al., 1997, Biswas et al., 2007) and shoot tip (Manjula et al., 1997, Soniya and Sujitha, 2006, Chandraprabha and Ramasubbu, 2010).

A survey of literature pertaining to the multiple shoot induction from the nodal and shoot tip explants in Aristolochiaceae revealed that addition of exogenous cytokinins is necessary before successful regeneration can occur. In *A.tagala*, BAP (3μM) along with KIN (0.5μΜ) was found to be the optimal combination for multiple shoots induction, indicating a synergestic effect of these two growth regulators. The synergestic effect of two cytokinins favouring adventitious shoot formation has also been demonstrated in the shoot tips of *Musa* spp. (Madhulatha et al., 2004), in the shoot tips and nodes of *Portulaca grandiflora* (Srivastava and Joshi 2009) and *Plecyphora aselliformis* (Santos-Diaz et al., 2003).

Polyphenolic exudation from the cultured explants which hampered normal culture establishment within 10 days of culture in *A. tagala* was controlled by the incorporation of 0.1% (w/v) activated charcoal. Activated charcoal was also effective in preventing phenolic exudation from the shoot tip and nodal cultures of *A. indica* (Manjula et al., 1997). Activated charcoal is a complex substance and the entire range of its effects on tissue culture media and the subsequent growth and morphogenesis of tissue cultures is unknown. It has been used in plant tissue culture media to improve culture growth and/or promote morphogenesis in a wide variety of species. Growth-promoting effects of activated charcoal have been attributed to adsorption of substances inhibitory to growth from the media produced either from breakdown of the media during autoclaving (Weatherhead et al., 1978) or by the cultures themselves (Fridborg et al., 1978).Earlier reports on the shoot tip (Chandraprabha and Ramasubbu, 2010) and nodal explants (Biswas et al., 2007) of *A.tagala* had no mention about the polyphenolic exudation. This could probably be because the region from which the plants were collected was different from that of the present study. This is yet another example of the environmental effect of gene expression.

In *A.tagala*, in order to ensure early elongation of the adventitious shoots developed *in vitro*, it was necessary to transfer the same to MS basal medium supplemented with GA_3_. A similar effect of the hormone was also reported by (Manjula et al., 1997) in *A. indica*. Although BAP stimulates multiple shoot formation it interferes with development and elongation of shoots (Wright et al., 1986). Dayal et al. 2003 reported that in *Cajanus cajan* a longer exposure to a combination of BAP and KIN was detrimental for further elongation of shoot buds and they had to be exposed to GA_3_ for their further development. In the present study, a better elongation of shoots was observed in liquid MS medium supplemented with GA3 than in solid medium. This may be due to low oxygen concentration in solid agar medium when compared to liquid medium as suggested by Newell et al., 2003.

In *A. tagala*, a combination of IAA, KIN and BAP was found to be significant in enhancing the length of the roots. This is in concurrence with the results obtained by Remeshree et al. 1994 in *A. bracteolata*.

In view of the various therapeutic claims of the leaves of *A.tagala*, an attempt to analyse the various bioactive components in the leaves, *in vitro* derived leaves and leaf derived callus of *A.tagala* has been made in the present investigation. Also, we have tried to find out if there is any change in the composition of bioactive components because of the culture conditions.

The current study revealed the presence of a wide variety of flavonoids in the extracts of *A.tagala*. Earlier reports indicated significant activity of flavonoids against several microorganisms (Bratner et al., 1996, Wang et al., 1988, Ismail and Alam, 2001). The presence of flavanoids gives credence to the use of *A.tagala* in treating fever and dysentery.

Phenols like gentisic acid, pyragallol, 2-methyl resorcinol, orcinol, catechol and sugars like ribose, rhamnose, galactose, and glucose were identified in the *ex vitro* leaves, *in vitro* leaves and leaf derived callus of *A. tagala*.. Naturally occurring phenolic compounds have free radical scavenging properties, due to their hydroxyl groups (Diplock, 1997). Further, phenolic compounds are effective hydrogen donors, which make them antioxidant (Rice Evans et al.,1995). The result obtained is compatible with one of the uses of *A.tagala* preparation in traditional medicine for treatment of rheumatism, since free radicals usually induce cellular damage and play a crucial role in many diseases such as rheumatism, cancer, hepatic disorder and aging disease (Auroma, 1994).

The alkaloid, berberine was identified in the *ex vitro* and *in vitro* leaf samples of *A. tagala. In vitro* experimental and clinical results indicate that berberine as an excellent disinfectant for infective root canal deciduous teeth (Su, 1992).This validates the claim of *A. tagala’s* use in tooth ache.

GCMS analysis of the plant samples showed the presence of sesquiterpenes, which are a class of naturally occurring molecules that have demonstrated therapeutic potential in decreasing the progression of cancer (Modzelewska et al., 2005). The presence of a sesquiterpene hydrocarbon, Ishwarane, has been observed from the *in vivo* and *in vitro* leaves of *A.tagala*. This is the first report for the presence of this compound in *A.tagala*. The same compound has been reported earlier from *A. argentina* (Priestap et al., 2003) and *A.indica* (Govindachari et al.,1970).

In the present investigation, the phytochemical profiles of *in vitro* and *in vivo* plant samples varied, which may be due to the influence of cultural environment, organic and inorganic components in the medium and growth hormones. Working with tissue cultures of *Catharanthus roseus*, Zenk (1978) observed that growth hormones added to the medium strongly influenced the production of phytochemicals. Manipulation of the plant cell culture environment and media can affect the rates of both cell growth and accumulation of secondary metabolites (Bhalsing and Maheshwari, 1998, Butcher, 1998).

## Conclusions

The present study on *A.tagala* provides an efficient, rapid and reproducible protocol for the mass multiplication and conservation of this threatened medicinal plant. This is highly advantageous for the conservation of this species and may further benefit aims of secondary metabolite isolation and genetic modification in *A.tagala*. This is the first report of analysis of bioactive compounds during the *in vitro* developmental stages of *A.tagala*. The dire need of the hour is to explore the potential metabolites of plant origin, which can mimic the effect of present drugs or can be supplemented with other drugs to make them more effective and easily available to mankind.

Competing interest

The authors declare that there is no competing interest.

## REFERENCES

Adams, R.P. (2001). Identification of Essential Oil Components by Gas chromatography/Mass Spectroscopy. Carol Stream, Illinois, US: Allured Publishing.

Archana, S. and Aruna, G.J. (2009). *In vitro* behaviour of nodal explants of *Portulaca grandiflora* under the influence of cytokinins. Acta Universitatis Latviensis. 753, 43–48.

Auroma, O.I. (1994). Nutritional and health aspects of free radical and antioxidants. Food and Chemical Toxicology. 32, 671–678.

Bate-Smith, E.C. (1962). Attractants and repellents in higher animals. In Phytochemical Ecology (ed. J.B. Harborne), pp. 45–46. Academic Press.

Banziger, H. and Disney, R.H.L. (2006). Scuttle flies (Diptera: Phoridae) imprisoned by *Aristolochia baenzigeri* (Aristolochiaceae) in Thailand. Mitteilungen der Schweizerischen entomologischen Gesellschaft. 79, 29–31.

Bhalsingh, S.R. and Maheshwari, V.L. (1998). Plant tissue culture - A potential source of medicinal compounds. Journal of Scienctific and Industrial Research. 57, 703–708.

Biswas, A., Bari, M.A., Mohashweta, R. and Bhadra, S.K. (2007). *In vitro* Regeneration of *Aristolochia tagala* Champ. A Rare Medicinal Plant of Chittagong Hill Tracts. Journal of Biological Sciences. 15, 63–67.

Bratner, A., Males, Z., Pepeljak, S. and Antolic, A.(1996). Antibacterial activity of *Paliurus spina-christi* Mill (Christ’s thorn). Journal of Ethnopharmacology. 52, 119–122.

Butcher, D.N. (1998). Secondary products in tissue cultures. In Plant Cell, Tissue, and Organ Culture (ed. J. Reinert and Y.P.S. Bajaj), pp.668–693. New Delhi, India: Narosa Publishing House.

Chandraprabha, A. and Rama Subbu, R. (2010). Micropropagation of *Aristolochia tagala* Cham. – A Rare and Endemic Medicinal plant from Western Ghats. Journal of biosciences research. 1, 70–73.

Chang, C.C., Yang, M.H., Wen, H.M. and Chern. J.C. (2002). Estimation of total flavonoid content in propolis by two complementary colorimetric methods. J. Food Drug Anal. 10, 178–182.

Dayal, S., Lavanya, M., Devi, P. and Sharma, K.K. (2003). An efficient protocol for shoot regeneration and genetic transformation of pigeon pea [*Cajanus cajan* (L) Mill sp.] using leaf explants. Plant Cell Reports. 21, 1072–1079.

Diplock, A.T. (1997). Will the ‘good fairies’ please prove to us that vitamin E lessens human degenerative disease? Free Radicle Research. 27, 511–532.

Dubois, M.K.A., Gilles, J.K., Hamilton, P.K. and Repers, F.(1956). Colorimetric method for determination of sugars and related substances. Anal. Chem. 28, 350-356

Fazel Shamsa., Hamidreza Monsef., Rouhollah Ghamooshi. and Mohammadreza Verdian-rizi. (2008). Spectrophotometric determination of total alkaloids in some Iranian medicinal plants. Thai J. Pharm. Sci. 32, 17–20.

Fridborg, G., Pedersen, M., Landstron, L.E. and Erikson, T. (1978). The effect of activated charcoal on tissue culture: adsorption of meatabolites inhibiting morphogenesis. Physiologia Plantarum. 43, 104–106.

Govindachari, T.R., Mohamed, P.A. and Parthasarathy, P.C. (1970). Ishwarane and Aristolochene, two new sesquiterpene hydrocarbons from Aristolochia indica. Tetrahedron. 26, 615.

Harborne, J.B. (1960). Phytochemical methods: A guide to modern technique of plant analysis. London: Chapman and Hall.

Harborne, J.B. (1973 a). Phytochemical methods: A guide to modern technique of plant analysis. London: Chapman and Hall.

Harborne, J.B. (1973 b). Phytochemical methods: A guide to modern technique of plant analysis. London: Chapman and Hall.

Ismail, N., Alam, M.(2001). A novel cytotoxic flavonoids glycoside from Physalis angulata. Fitoterapia. 72, 676–679.

Klitzke, C.F. and Brown, K.S. (2000). The occurrence of aristolochic acids in neotropical troidine swallowtails (Lepidoptera: Papilionidae). Chemecology. 10, 99–102.

Kritikar, K.R. and Basu, B.D. (2003). Indian medicinal plants, Volume II. India: International book distributors.

Kumar, V., Poonam Prasad, A.K. and Parmar, V.S. (2003). Naturally occurring aristolactams, aristolochic acids and dioxoaporphines and their biological activities. Natural Products Report. 20, 565–583.

Madhulatha, P., Anbalagan, M., Jayachandran, S. and Sakthivel, N. (2004). Influence of liquid pulse treatment with growth regulators on *in vitro* propagation of banana (*Musa* sp. AAA). Plant Cell Tissue and Organ Culture. 76, 189–191.

Manjula, S., Thomas, A., Daniel, B. and Nair, G.M. (1997). *In vitro* plant regeneration of Aristolochia indica. Phytomorphology. 47, 161–165.

McDonald, S., Prenzler, P.D., Antolovich, M. and Robards, K.(2001). Phenolic content and antioxidant activity of olive extracts. Food Chem. 73, 73–84.

Modzelewska, A., Sur, S., Kumar, S.K. and Khan, S.R. (2005). Sesquiterpenes: natural products that decrease cancer growth. Current Medicinal Chemistry. Anticancer Agents. 5, 477–499.

Murashige, T. and Skoog, F.(1962). A revised medium for rapid growth and bioassays with tobacco tissue culture. Physiologia Plantaraum. 15, 473–497.

Murugan, R., Shivanna, K.R. and Rao, R.R. (2006). Pollination biology of *Aristolochia tagala*, a rare species of medicinal importance. Current Science. 91, 795–798.

Newell, C., Growns, D. and Mc Comb, J. (2003). The influence of medium aeration on *in vitro* rooting of Australian plant microcuttings. Plant Cell Tissue and Organ Culture. 75, 131–142.

Priestap, A.H., Van Baren, C.M., Lira, P.D.L., Coussio, J.D. and Bandoni, A.L. (2003). Volatile constituents of *Aristolochia argentina*. Phytochemistry. 63, 221–225.

Ravikumar, K. and Ved, D.K. (2000). 100 Red Listed Medicinal Plants of Conservation Concern in Southern India. Bangalore, India: Foundation for Revitalization of Local Health Traditions.

Remeshree, A.B., Hariharan, M. and Unnikrishnan, K. (1997). In vitro organogenesis in *Aristolochia indica*. Phytomorphology. 47, 161–165.

Remeshree, A.B., Hariharan, M. and Unnikrishanan, K. (1994). Micropropagation and callus induction of Aristolochia bracteolate Lam. a medicinal plant. Phytomorphology 44, 247–252.

Remya, M., Narmatha Bai, V. and Mutharaian, V.N. (2013). In vitro regeneration of Aristolochia tagala and production of artificial seeds. Biologia Plantarum. 57, 210-218.

Rice-Evans, C.A., Miller, N.J., Bolwell, P.G., Bramley, P.M. and Pridham, J.B. (1995). The relative antioxidant activity of plant derived polyphenolic flavonoids. Free Radicle Research. 22, 375–383.

Santos-Diaz, M.A., Mendez-Ontiveros, R., Arrendono-Gomez, A. and Santos-Diaz, M.D.L. (2003). *In vitro* organogenesis of *Pelecyphora aselliformis* Eshenberg (Cactaceae). In vitro Cell and Devlopmental Biology. 39, 480–484.

Soniya, E.V. and Sujitha, M. (2006). An efficient in vitro propagation of *Aristolochia indica*. Biologia Plantarum. 50, 272–274.

Su, Y.H., Liu, Y.B. and Zhang, X.S. (2011). Auxin–Cytokinin Interaction Regulates Meristem Development. Molecular Plant. 4, 1–11.

Trujillo, C. and Sersic, A. (2006). Floral Biology of *Aristolochia argentina* (Aristolochiaceae). Flora. 201, 374–382.

Wang, F.M. and Xueyl (1988). The changes in polyamine content and polyamine oxidase activity during in vitro bulblet formation from bulb scales of *Lilium davidii* var. Unicolor. Acta Phytophysiol. Sinica. 14, 350–355.

Weatherhead, M.A., Burdon, L. and Henshaw, G.G. (1978). Some effects of activated charcoal as an additive to plant tissue culture media. Z. Planzenphysiol. 89, 141–147.

Wright, M.S., Carner, M.G., Hinchee, M.A., Davis, G.C., Koehler, S.M., Williams, M.H., Colburn, S.M. and Pierson, P.E. (1986). Plant regeneration from tissue cultures of soybean by organogenesis. In Cell culture and somatic cell genetics of plants (ed. W.H. Parrot and F.L. White), pp. 111–119. London: Academic press.

Zenk, M.H. (1978). The impact of plant cell culture on Industry. In Frontiers of Plant tissue Culture, Proc. 4^th^ Intl. Congress Plant Tiss. Cell Cult. (ed. T.A. Thrope), pp.1–13. Calgary, Canada.

